# Exogenous supply of Hsp47 triggers fibrillar collagen deposition in skin cell cultures *in vitro*

**DOI:** 10.1101/803791

**Authors:** Essak S Khan, Shrikrishnan Sankaran, Lorena LLontop, Aránzazu del Campo

**Affiliations:** INM – Leibniz Institute for New Materials, Campus D2 2, 66123 Saarbrücken, Germany; Chemistry Department, Saarland University, 66123 Saarbrücken, Germany

**Keywords:** Hsp47, collagen defect, skin, 3-fold increase, firbrillar collagen

## Abstract

**Background:** Collagen is a structural protein that provides mechanical stability and defined architectures to skin. In collagen-based skin disorders like Epidermolysis bullosa, EDS the ability to offer such stability is lost either due to mutations in collagens or defect in the chaperones involved in collagen assembly, which leads to chronic wounds, skin fragility, and blisters. Existing approaches to study and develop therapy against such conditions are the use of small molecules like 4-phenylbutyrate (4-PBA) or growth factors like TGF-β. However, these approaches are not collagen specific resulting in unsolicited responses. Therefore, a collagen specific booster is required to guide the correct folding and deposition of collagen in a highly regulated manner. Hsp47 is a chaperone with a major role in collagen biosynthesis. Expression levels of Hsp47 correlate with collagen production. This article explores the stimulation of collagen deposition by exogenously supplied Hsp47 (collagen specific chaperone) in skin cells, including specific collagen subtypes quantification.

**Results:** Here we quantify the collagen deposition level and the type of deposited collagens by different cell types from skin tissue (fibroblasts NHDF, L929 and MEF, keratinocytes HaCat and endothelial cells HDMEC) after Hsp47 stimulation. We find upregulated deposition of fibrillar collagen subtypes I, III and V after Hsp47 delivery. Network collagen IV deposition was enhanced in HaCat and HDMECs and fibril-associated collagen XII were not affected by the increased Hsp47 intracellular levels. The deposition levels of fibrillar collagen were cell-dependent i.e. Hsp47-stimulated fibroblasts deposited significantly higher amount of fibrillar collagen than Hsp47-stimulated HaCat and HDMECs.

**Conclusions:** A 3-fold enhancement of collagen deposition was observed in fibroblasts upon repeated dosage of Hsp47 within the first 6 days of culture. Our results provide fundamental understanding towards the idea of using Hsp47 as therapeutic protein to treat collagen disorders.

## Introduction

Collagen fibers represent 60-80% of skin dry weight and confer skin its resistance to mechanical stress [1–4]. The skin is a layered tissue, and the collagen composition and morphology of each layer is different [5, 6]. COL I is predominant in the dermal and hypodermal layer, and forms heterotypic structures with other collagens such as COL III and/or V [7]. The basement membrane separating the epidermis and dermis is rich in COL IV. In multiple skin pathologies collagen organization is altered, either genetically or acquired due to environmental factors. Genetic collagen-related skin disorders such as Epidermolysis bullosa (EB) [8] and Ehlers-Danlos Syndrome (EDS) are both caused due to mutations in fibrillar COL I[9] and/or COL III[10]. The patients have fragile skin, blisters and chronic wounds as consequence of reduced collagen levels in the skin tissue due to collagen miss folding, impaired formation of highly organized structures, poor collagen crosslinking, and accelerated collagen degradation[11]. Scurvy and Aging have localized wrinkles and blisters due to weakening of skin structural architecture between dermis and epidermis due to sparse collagen fiber density and extensive degradation of fibrillar collagen, mostly COL I [12, 13] by matrix metalloproteinase [14, 15]. The existing therapies for these disorders are based on the delivery of growth factors (e.g. TGF-beta[16, 17]) and chemical stimulants (e.g. ascorbic acid [17–19], glycolic acid[20], 4-phenyl butyric acid (4-PBA)[21] and retinol[22]) to boost the collagen production and matrix deposition. However, these molecules have multiple other roles in the body and the therapies are associated with negative side effects, such as promoting abnormal angiogenesis, or inflammatory responses.

We recently demonstrated that uptake of exogenous Hsp47 could specifically enhance collagen deposition in fibroblast cultures[23]. Uniquely, Hsp47 is a collagen-specific chaperone. It has multiple roles in collagen biosynthesis, i.e. it stabilizes triple helical of procollagen[24–28], it prevents intracellular procollagen degradation[29–31], it is involved in quality control of folded procollagen [31, 32], it inhibits procollagen aggregate formation in the Endoplasmic Reticulum (ER) [33, 34], and it supports procollagen transport to Golgi apparatus[30] by binding to procollagen in the ER (at neutral pH) and dissociating in the cis-Golgi (at low pH). The involvement of endogenous Hsp47 in the biosynthesis of collagen subtypes I to V has been reported [23, 29, 35, 36]. It is however unclear if exogenous administration of Hsp47 to cells affects the deposition of each collagen subtype to a similar level, or if deposition of certain collagen is preferentially supported. Note that expression levels of Hsp47 are altered in some variants of EB[37] and EDS[29] and this protein is up regulated in cancer[38–40].

Hsp47 is retained in the ER via KDEL receptor mediated transport from the Golgi to the ER [26, 29, 41–45]. This KDEL-receptor is also found on the cell membrane,[46] and is critical for the transport of exogenous Hsp47 to the ER of cells *in vitro* by simple incubation with the protein in solution[23]. Expression levels of KDEL receptor at the cell membrane are cell-dependent [47, 48]. Therefore, Hsp47 uptake and the resulting up regulated collagen expression levels by exogenous Hsp47 might also be cell-dependent.

In the current work we follow-up on our previous demonstration of Hsp47-stimulated deposition of collagen in *in vitro* in fibroblast cultures[23], and we quantify the cell- and type-specific collagen deposition (fibrillar collagens I, III, V, network collagen IV, and fibril-associated collagen XII) after Hsp47 stimulation in *in vitro* cultures of fibroblasts, epithelial and endothelial cells from skin tissue. Lastly, the increase in the collagen deposition upon repetitive delivery of recombinant Hsp47 was also studied. Our results provide fundamental understanding of the potential of Hsp47 as therapeutic protein.

## Results

The uptake of Hsp47 was tested in fibroblast (NHDF from human skin dermis, L929 from mouse adipose tissue and MEF from mouse embryos), epithelial (HaCaT, human epidermal keratinocytes) and endothelial (HDMEC from human dermis) skin cells. A recombinant EGFP-tagged Hsp47 (hereafter referred to as H_47[23]_) was used for these studies. The different cell types were seeded at same cell density and incubated with H_47_ at concentrations between 0.1 μM and 1.0 μM. Incubation with lower H_47_ concentrations showed no detectable fluorescence signal inside the cells. Cells incubated with >1 μM H_47_ showed fluorescent aggregates on the surface of the culture plate, indicating that saturation levels of H_47_ for uptake were achieved and H_47_ was binding to extracellular collagen deposited by the cells. At the intermediate concentrations accumulation of green fluorescence was detected inside the cells and no aggregates were observed outside of the cells, indicating efficient uptake of H_47_. The incubation time of 3 h was uptaken from our previous experience[23].

H_47_ uptake was visualized by epifluorescence imaging of EGFP green signal inside the cells. Co-localization of H_47_ and the ER signals confirmed accumulation of the uptaken H_47_ at the ER in a cell types (Figure 2b, S1). No fluorescence signal was observed when cells were incubated with EGFP alone, indicating that uptake is specific to the H_47_ sequence and not mediated by the EGFP label (Figure S1 and 2). Our previous work demonstrated that this occurs through KDEL receptor mediated endocytosis and transport[23]. H_47_ uptake was quantified by counting the percentage of cells with detectable ER- and H_47_ fluorescent signals after incubation for 3 h and medium change. 100% uptake indicates that all the cells with labeled ER contained labeled H47. Results show that H_47_ uptake was concentration dependent, and cell type dependent (Figure 2a). An increase in the H_47_ incubation concentration led to increased uptake, and >80% uptake was observed for fibroblasts and endothelial cells at 0.5 μM H47. Saturation levels are achieved at 0.5 and 0.8 μM H_47_ for MEF and L929 cells, and at 1 μM H_47_ for NHDF and HDMECs. HaCaT cells showed the lowest uptake, with maximum uptake values of ca. 80% achieved at >0.8 μM H_47_ concentrations. This variation is attributed to differences in distribution and density of KDEL receptors on the cell membrane, which has been reported by other authors[46].

**Figure 1.**
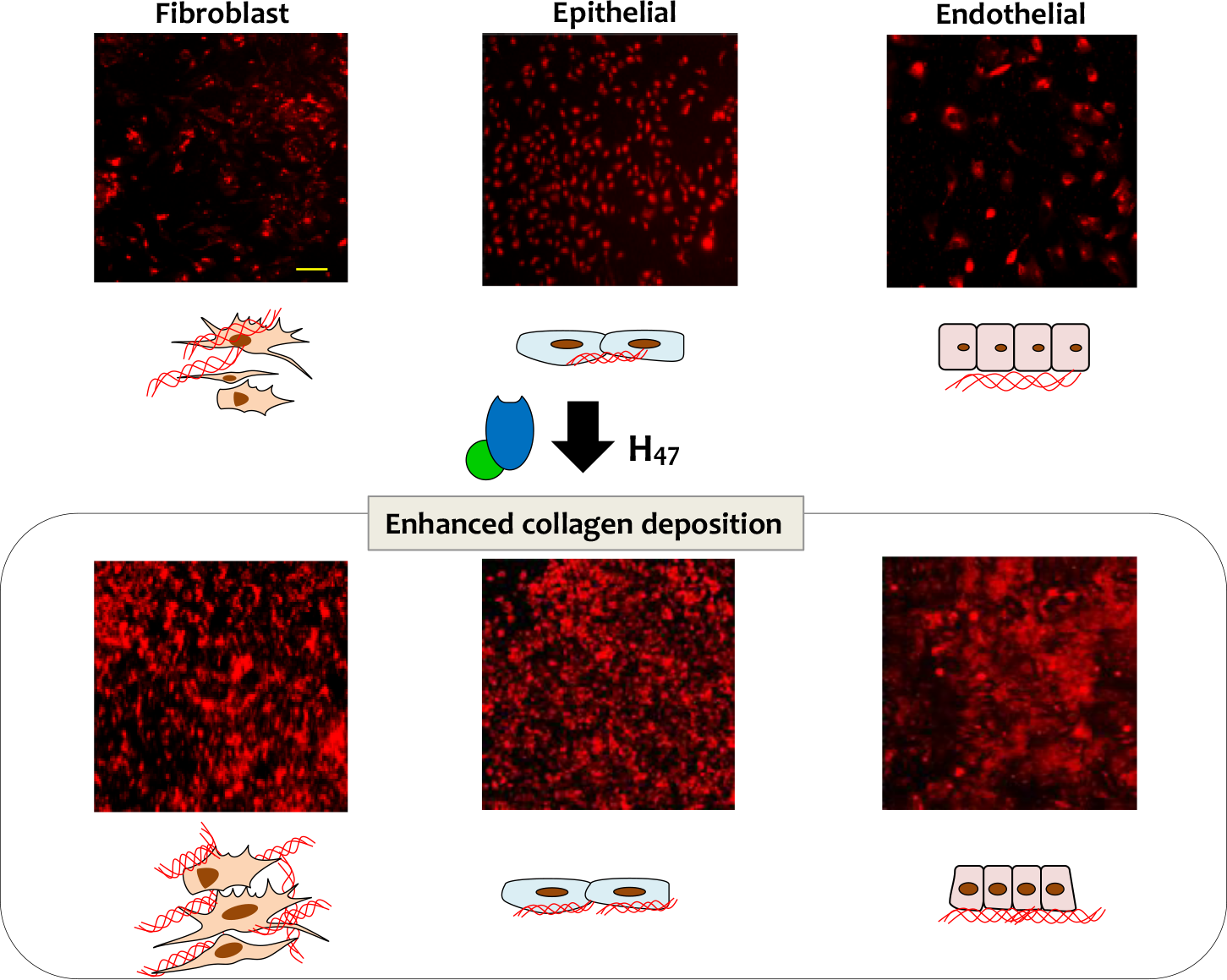
Scheme showing enhanced collagen deposition by treatment with recombinant H47. Immuno-staining of COL I in fibroblast, epithelial and endothelial cell lines is shown in Red using COL I antibody. Scale: 250μm.

**Figure 2.**
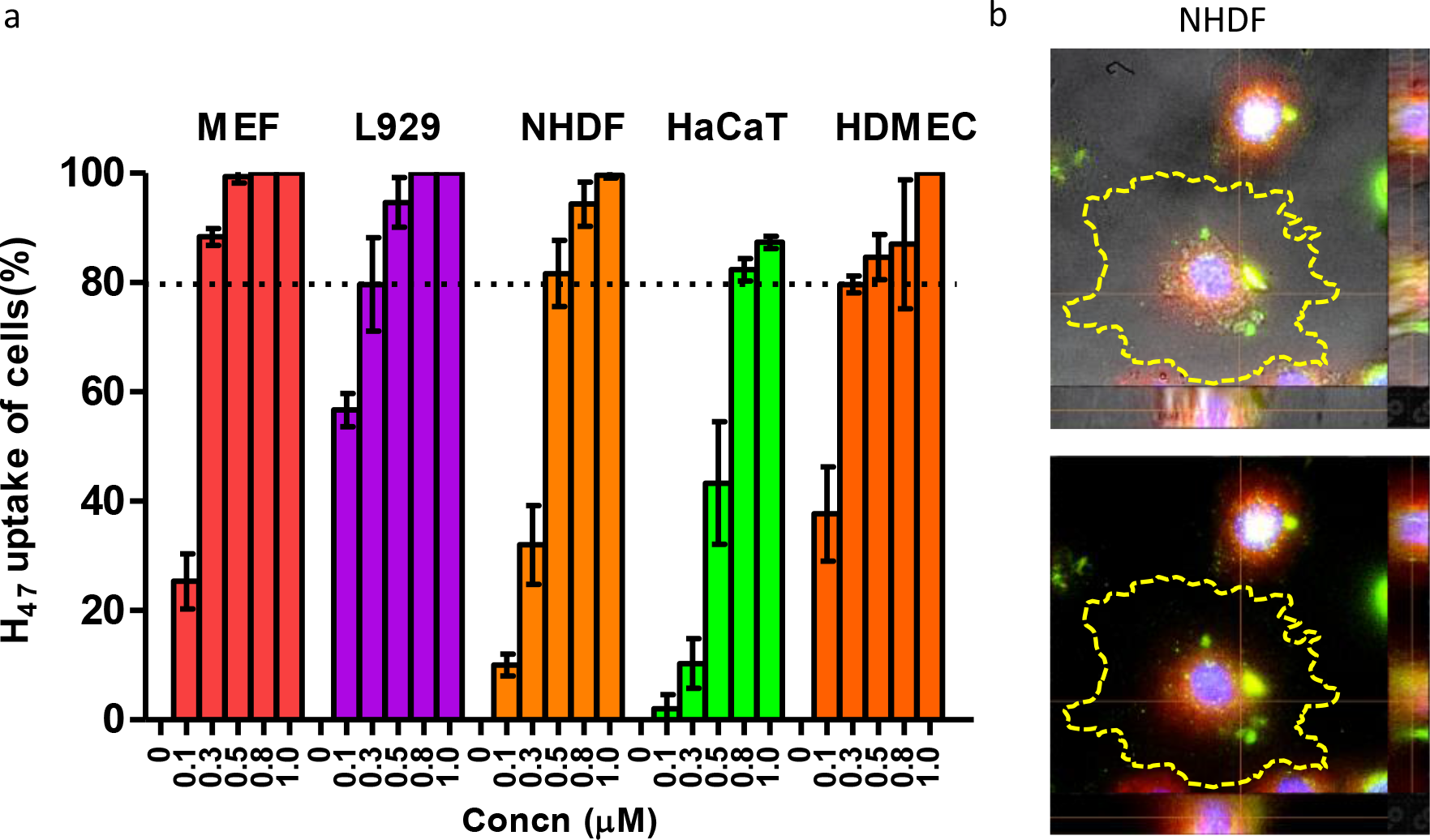
**a.** H47 uptake by NHDF, L929, MEF, HaCaT and HDMEC. Cells were incubated with increasing concentrations of H_47_ in the medium (0.1μM-1.0μM) for 3 h. The % of H_47_ uptake was obtained by quantifying the percentage of cells showing H_47_ (EGFP green signal) and ER tracker dye (Red signal). 100% H_47_ uptake of cells indicate that all cells in culture had co-localized signal. Error bars indicate standard deviation of n=3 experiments. **b**. Z-stack fluorescence images showing up taken H_47_ (green) co-localized with ER (red) in NHDF (Blue: DAPI). The yellow dotted line around the cells indicate the edge of the cells. Scale: 20 μm

We then compared H47-induced collagen deposition by the different cell types after incubation with 0.5 μM H_47_. For this purpose, cells were seeded for 24 h on tissue culture plastic wells and incubated with 0.5 μM of H_47_ for 3h. The medium was exchanged and cells were cultured for further 24h. The deposited collagen on the culture plate was labeled with Picro Sirius Red and quantified by spectrophotometry. Sirius red is a strong anionic dye comprising six sulfonate groups that binds preferentially to the cationic groups of the collagen fibers [49, 50]. Data were normalized by the value of collagen deposition by MEF cells without any treatment. The fibroblast cell lines MEF, L929 and NHDF showed a 70% −100% increases in collagen deposition when treated with H_47_. Under the same incubation conditions, the epithelial (HaCaT) and endothelial (HDMEC) cell lines showed 20 and 50% increase in collagen deposition respectively (Figure 3a). These results indicate that exogenously supplied H_47_ induces collagen deposition more effectively in fibroblast cells. This is in agreement with the natural role of fibroblasts as major matrix-producing cells in connective tissue [51, 52].

**Figure 3.**
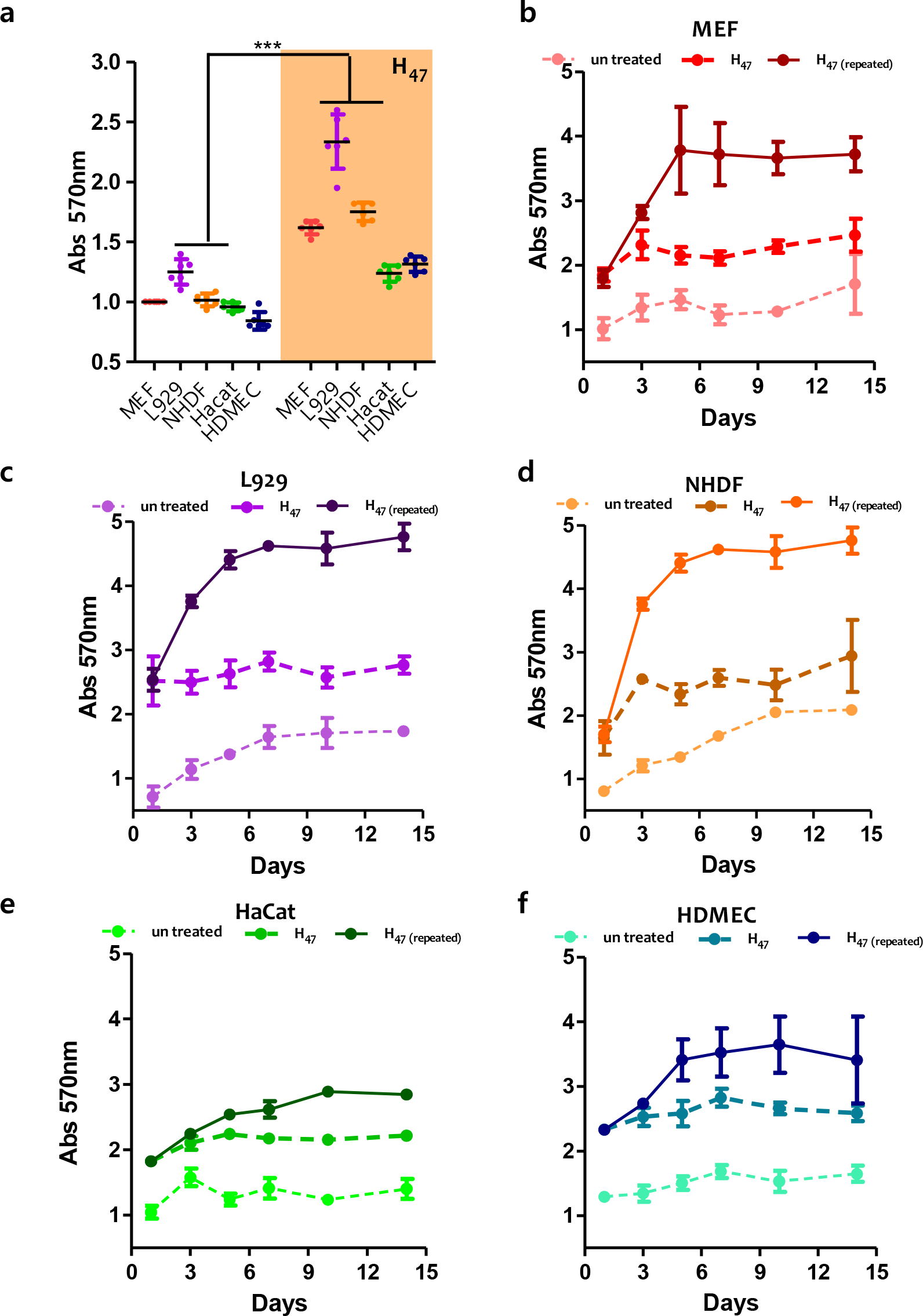
**a.** Quantification of collagen deposition using Sirius Red assay in different cell types 24h after H_47_ treatment. The plots were normalized by untreated MEF condition. Statistical significance for both b and c was analyzed by Tukey test comparing untreated against H_47_ treated cells (mean±SD, ANOVA, *** *p*<0.001). Error bars representing standard deviation of 6 independent experiments. **b-f**. Quantification of collagen deposition using Picro Sirius Red assay in **b.** MEF, **c.** L929, **d.** NHDF, **e.** HaCat and **f.** HDMEC cells on days 1,3,5,7,10 and 14. Error bars represent standard deviation from 3 individual experiments.

We followed collagen deposition at increasing culture times up to 14 days. Collagen deposition in Fibroblasts (L929, NHDF, MEF) without H_47_ treatment increased during the first 6 days and reached a steady state at longer time scales (Figure 3 b, c and d). This trend correlates with proliferation kinetics expected for this cells, and the expected down regulation of matrix production when confluency is reached[53]. Fibroblast cells were 80-100% confluent at day 9-10. HaCaT and HDMEC cells did not show an appreciable increase in collagen deposition with time, and the overall amount of deposited collagen was much lower (Figure 3 e and f). When cultures were treated with H_47_ on day 1, the total amount of deposited collagen at least doubled on day 1, but the collagen deposition profile at longer times did not change significantly, i.e. the curve was just shifted to higher collagen values from day 1 (Figure 3). This result is in agreement with the expected life time of H_47_ (more than 24h for natural Hsp47 at physiological conditions [23, 54]). In addition, these results indicate that the collagen deposited after H_47_ treatment has similar stability to the naturally deposited collagen without H_47_ stimulation.

We then tested if the amount of deposited collagen could be further increased by repetitive delivery of H_47_ on consecutive days. Culture medium was supplemented with 0.5 μm H_47_ on days 1, 3, 6, 9 and 12, and collagen deposition was quantified on days 1, 3, 5, 7, 10 and 14. A 2-3fold increase in collagen deposition was observed in NHDF, MEF and L929 cells after repeated treatment with H_47_ until day 5. HaCaT and HDMEC cells showed 1-1.5fold increase in collagen deposition within the same timescale. Addition of H_47_ on days 6, 9 and 12 did not result in further changes in the deposited amount of collagen. This plateau of collagen deposition also occurred in the control experiments, in which cells were not treated with H_47_ within the same period of time, and is again associated with the achievement of confluency in the 2D cell culture. Confluency results in cell senescence, up-regulation of MMPs (especially MMP1) and down regulation of MMP inhibitors and ECM proteins like collagens and elastin at genetic levels [55–58]. In order to confirm this hypothesis, H_47_ treatment was performed on 2D L929 cell cultures once they reached confluence (Day 10). No significant increase in collagen deposition was observed 24 h after this treatment (Figure. S5b), corroborating our hypothesis. In addition, H_47_ was found bound to matrix collagen on the culture substrate, with no detectable delivery inside the cells (Figure. S5a). In summary, repeated dosage of Hsp47 increased collagen production in 2D cultures of skin cells until cells reached confluency.

The composition of the collagen matrix is tissue-dependent and different cells are expected to produce different collagen types. We investigated if Hsp47-induced collagen had the same composition as the naturally secreted collagen. For this purpose, H47-treated cultures were decellularized and the remaining matrix layer on the culture plate was stained using antibodies specific for COL I, III, V and XII. These collagens were selected based on their abundance in skin tissue and their involvement in skin related disorders, in which they are reduced or mutated [9–11, 22, 52]. The relative abundance of each collagen subtype was obtained from the fluorescence image of the culture plate. The mean fluorescence intensity value for each collagen subtype was corrected by subtraction of the background (see experimental details), and normalized by the corresponding value obtained in untreated MEF cells. An increase in the deposition of fibrillar COLs I, III and V was observed in all cell types upon H_47_ treatment. The increase in the deposition of fibrillar collagens I, III and V was significantly higher in fibroblast cultures *vs.* HaCaT and HDMEC cultures. Conversely, deposition of network COL IV upon H_47_ treatment was only enhanced on HaCaT and HDMECs (Figure 4 and S3). No changes were observed in the deposition of COL XII, suggesting that Hsp47 may not be involved in the secretion of the fibril-associated COL XII. We observed that cell spreading increased in cultures treated with Hsp47, which is an expected finding as collagen is a matrix protein with multiple adhesion sites for the integrin family.

**Figure 4.**
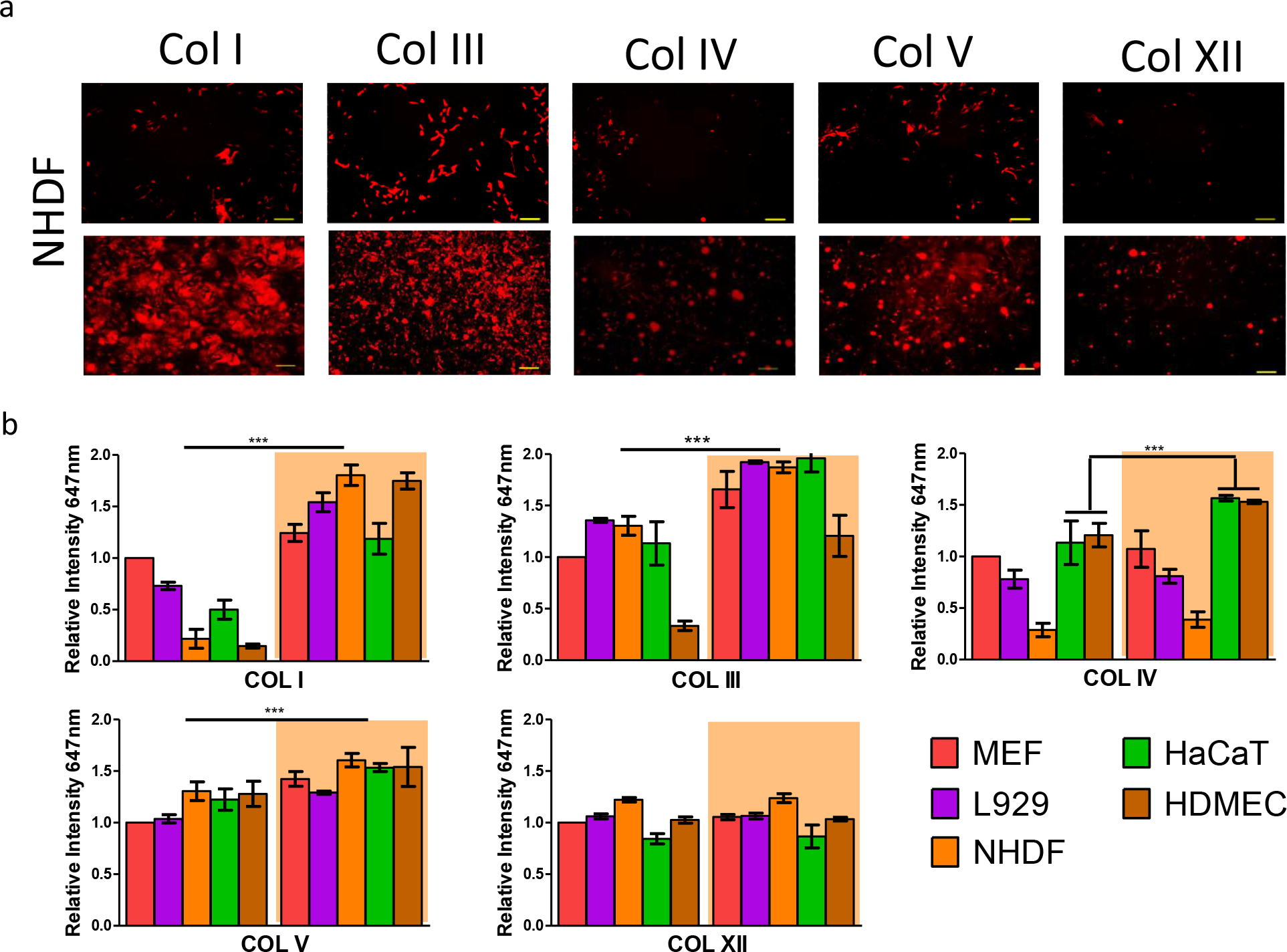
Quantification of H_47_ stimulated deposition of different collagen subtypes. **a.** Fluorescence images showing immunostained COL I, III, IV, V and XII in NHDF cells before and after treatment with H_47_ (Scale – 250μm) **b.** Quantification of deposited COL I, III, IV, V and XII from immuno-stained images in NHDF, L929, MEF, HaCaT and HDMEC cultures. Data correspond to collagen deposition 24 hours after H_47_ treatment and controls. The plots were normalized with MEF cells untreated condition as 1. Error bars represent standard deviation from n-3 experiments. Statistical significance was analyzed by Tukey test comparing non treated against H_47_ treated cells (mean±SD, ANOVA, *** *p*<0.001).

We also studied the influence of exogenously supplied H_47_ on collagen production in a previously established MEF Hsp47-knockout fibroblast cell line (Hsp47−/−), which does not produce endogenous Hsp47 [59–61]. Deposition of fibrillar COLs I, III and V increased upon H_47_ treatment. No increase in COL IV and XII was observed, in agreement with previous observations in the other fibroblast cell lines (Figure5 a, c, S4). In order to quantify deposited collagen subtypes at higher sensitivity, western blot analysis of the deposited matrix was performed. Higher deposition of COL I, III and V was confirmed, and a very low amount of COL IV was also detected. Interestingly, similar analysis performed with healthy MEF cells (Hsp47+/+) revealed that these cells produced these collagen types to similar extent as observed in Hsp47−/− cells after H_47_-induction (Figure 5b). These results demonstrate that treatment of H_47_ deficient cells with exogenous H_47_ restores the ability of the cells to deposit collagen at levels and composition similar to healthy cells. Taking into account the physiological relevance of matrix composition and properties in cellular behavior *in vivo* we speculate that treatment with Hsp47 could be a useful approach to enhance collagen synthesis and matrix deposition on-demand.

**Figure 5.**
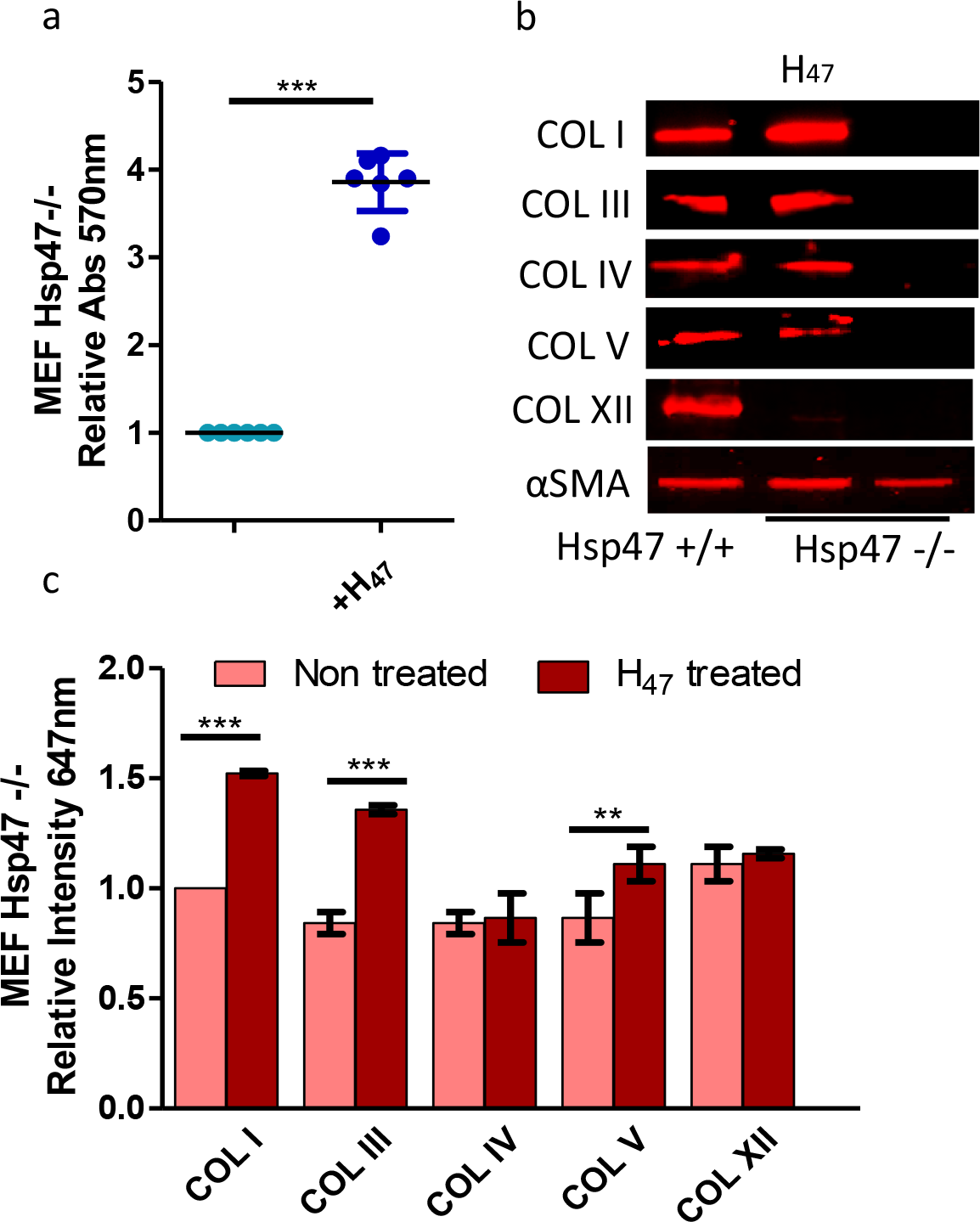
**a.** Quantification of fibrillar collagen deposited in MEF Hsp47-/* and Hsp47 +/+cells (control) at 24 hours after H_47_ treatment using Sirius Red assay. **b.** Western blot of COL I, III, IV, V and XII in deposited collagen from MEF Hsp47 +/+, Hsp47 −/* and Hsp47−/− cultures 24 hours after treatment with 0.5 μM H47. Red bands indicate signal from collagen subtypes specific antibody. **c**. Quantification of deposited COL I, III, IV, V and XII from immuno staining assays in MEF Hsp47−/− cultures. Error bars representing standard deviation from n-3 experiments in **a** and **c**, The plots in both **a** and **c** assays were normalized by MEF Hsp47−/− cells untreated condition taken as 1. Statistical significance in **a** and **c** was analyzed by Tukey test. Significance was calculated by comparing non treated against Hsp47 treated cells (mean±SD, ANOVA, *** *p*<0.001

## Discussion

Hsp47 is a collagen-specific chaperone protein with multiple roles in collagen biosynthesis. Expression levels of Hsp47 correlate with collagen production [59, 62]. Hsp47 has a regulatory role during scar formation in neonatal mouse skin after injury, which indicates that its expression during healing is up regulated *in situ*, in response to injury [63]. These facts indicate that Hsp47 could be an interesting therapeutic target in collagen-related skin disorders, as alternative to non-specific collagen inducers such as TGF β[64], VEGF[65] or ascorbic acid[17, 18]. Therapeutic use of these molecules to enhance collagen deposition influences other cellular functions such as proliferation[17, 18, 66, 67], differentiation[66] and angiogenesis[68], leading to undesired side effects. For example, ascorbate increases collagen production by acting as a co-factor to proline and lysine hydroxylases, which are involved in the hydroxylation of procollagen[42]. However, these enzymes are also involved in the hydroxylation of other matrix proteins, like Elastin or Fibronectin[69]. In contrast, the unique collagen-specificity of Hsp47, demonstrated in the literature [23, 29, 49], would allow up regulation of collagen deposition, without affecting any other molecule or cellular pathway.

The deposition of fibrillar collagens I and II, and to a less extent fibrillar collagen V was mainly enhanced in all tested cell types. Deposition of network COL IV was only observed in cells that are naturally connected to a basement membrane, where COL IV is also a major component. Fibril-associated COL XII was not enhanced by H_47_ treatment. The significant enhancement of fibrillar collagen deposition vs. other collagen types highlights the supporting role of Hsp47 in the intracellular assembly, stabilization and transport of collagen superstructures. A fibrillar collagen specific stimulation of collagen production could be a positive aspect for a potential Hsp47-derived therapy to enhance natural collagen production in diseases.

It is interesting to compare the efficiency of Hsp47-based enhancement in collagen deposition vs. treatment with other collagen inducers from literature data. In the comparison both the amount of deposited collagen and the time scale at which noticeable deposition occurred is relevant. We anticipate that the comparison is done among different cell types and culture methods and, therefore, numbers can only be taken as indicative. In our data, the deposition of fibrillar COLs I and III in fibroblasts (MEF, L929 or NHDF) was enhanced up to 70% −100% in 24 h. In contrast, ascorbate treatment of primary healthy skin fibroblast increased collagen production by 20-40 % after 4 days of treatment *in vitro* (measured by radioactively labeling procollagen),[70] and only 10 % in Hsp47 deficient fibroblast (measured by Sirus red assay)[23]. Glycolic acid treatment increased collagen production by 48% in a week in human skin fibroblast culture from neonatal foreskin and outgrown cells [71] and Vitamin A (retinol) induced a 100% increase in chronological aging skin in human patients *in vivo* after 24 weeks[72]. This comparison reveals that Hsp47 promotes fast and efficient deposition of fibrillar collagen in comparison with the other molecules, which would be a beneficial feature for a therapeutic use of Hsp47.

*In vivo* collagen assembles into high order structures[73]. In 2D cultures the interaction of the secreted collagen with the polystyrene plate affects the assembly [74]. Studies in real tissues would be necessary to proof if the secreted collagen upon Hsp47 treatment is able to form morphologically complex structures as it occurs in the natural extracellular matrix.

## Conclusions

The collagen subtype distribution in natural tissue plays a crucial role in tissue biomechanics, and alterations result in pathological states. In the skin, alterations in collagen levels are present in EDS, EB or Scurvy, and lead to chronic wounds, blisters and skin fragility. Our results show that exogenous delivery of Hsp47 chaperone can enhance collagen deposition in a cell-specific manner. Being a collagen specific molecular chaperone, Hsp47 treatment is not expected to affect other cellular process, unlike other collagen-stimulators like ascorbic acid, glycolic acid, retinol or growth factors would do. This is a relevant advantage Hsp47 for its potential use as matrix-stimulating therapeutic protein.

## Materials and methods

### Synthesis and Purification of H_47_

EGFP-Hsp47 (H_47_) was synthesized and purified using previously established protocol[23]. Pre-inoculums of H_47_ were grown overnight in LB medium containing NaCl (5g/L) and kanamycin (Kan) (25μg/ml) at 37°C/250 rpm. After 24 h, the cultures were transferred into a 2 L flask and were grown until OD600 of 0.65 in 1L LB NaCl Kan media at 37°C/250 rpm for proper aeration. The protein expression was induced with 1.0 mM isopropyl-β-d-thio-galactoside and was incubated overnight at 30°C/180 rpm. The cultured cells were harvested and pellets were stored at −80°C. Stored cells thawed and were resuspended in lysis buffer (300 mM NaCl, 10 mM imidazole, 20 mM Tris, pH 8) and lysed by sonication. Cleared lysate was purified by Ni-NTA affinity chromatography (Ni-NTA superflow; Qiagen). The soluble fraction was concentrated and purified further by 50K Advanced Centrifugal Device (Macrosep) in buffer A [20 mM Hepes (pH 7.5), 300 mM NaCl, and 4 mM DTT].

### Cell delivery assay

Hsp47+/+ and Hsp47−/− MEFs derived from Lethal Mouse Embryos [61] were gifted by Prof. Dr. Kazuhiro Nagata, Kyoto Sangyo University, Japan, L929 fibroblasts (ATCC CRL-6364), NHDF(Promo Cell C-12302), HaCaT (ATCC® PCS-200-011) and HDMEC (Promo Cell C-12210) were purchased from commercial suppliers.

MEF, L929, NHDF and HaCaT cells were seeded on 15 well Ibidi μ-Slide Angiogenesis plates (20,000 cells per well) with DMEM GlutaMax (Gibco) containing 10% fetal bovine serum (FBS; Gibco) and 1% Penicillin-Streptomycin (Pen-strep) antibiotic. HDMEC cells (20,000 cells per well) were seeded using M199 medium with endothelial growth supplement (Gibco), 10% fetal bovine serum (FBS; Gibco) and Pen-strep antibiotic. After 24 h, cells were incubated with varying concentrations of H_47_ (0.1μM-1μM) for 3 h. After incubation, the medium was removed and cells were washed once with sterile Assay buffer (1X) provided in the kit. A Dual staining solution (for nucleus and ER) was prepared by mixing 1 μl of ER tracker dye (ER Staining Kit - Red Fluorescence - Cytopainter (ab139482)) with 1 μl of DAPI in 1 ml of ER Assay buffer (1X) provided in the kit. The cells were incubated with 60μl of Dual staining solution per well at 37ᵒC for 1 h. Cells were washed with Assay buffer (1X) once, fixed with 4% PFA for 10 mins and washed three times with Assay buffer (1X). All the experiments were done in triplicate.

### Quantification of H_47_ uptake by different cell types

Imaging of the cell cultures was performed using a Nikon Ti-Ecllipse microscope (Nikon Instruments Europe B.V., Germany) with a 60X objective. The number of cells showing both H_47_ signal with ER tracker signal were counted and represented in percentage. Maximum of 100% was considered if all cells showed EGFP green signal from H_47_ with ER signal. This analysis was performed on all cell types. All the analysis was done using Image J and graphs were plotted using graph prism software for three independent experiments for each cell types.

### Sirus Red Assay for quantification of Collagen deposition and Immuno-staining

Cells were cultured in 24 well plate for 24 hours (50K cells per well). Cells were incubated with 0.5 μM solutions of H_47_ in DMEM and M199 medium for 3 h, followed by medium exchange and cultured for 1, 3, 5, 7, 10 and 14 days. Non treated cells were cultured for similar time points and used as controls in this experiment. For testing repeated H_47_ treatments in all the cell types, H_47_ was also added on day 1, 3, 6, 9 and 12. Collagen deposition was quantified on days 1, 3, 5, 7, 10 and 14 by Sirus Red assay as mentioned above. For Sirius Red assay cells were fixed using Bouin solution (75% picric acid, 10% formalin, and 5% acetic acid) (Sigma HT10132). Collagen deposited in the wells was stained by incubating with 0.1% Sirius red in picric acid (ab150681) for 1 h and washing with 0.01 N HCl. The matrix was dissolved in 0.1 N NaOH and the absorption of the slurry were measured at 570 nm using a Biolumin960k spectrophotometer [23, 49]. The absorbance values (Figure 3a in manuscript) were normalized by the value of collagen deposition from untreated MEF cells in 24h. In MEF Hsp47 −/− untreated cells were considered as 1 for Figure 4a.

For testing ability of H_47_ to enhance collagen deposition on cells reaching confluency, L929 were seeded at a high density (50 cells/well) and allowed to become confluent within 2 days and treated with H_47_ and kept for 24h after treatment. Collagen deposition was quantified using sirus red assay as mentioned above. The absorbance values were normalized by the value of collagen deposition from untreated L929 cells in 24h as 1. Also deposited collagen was stained with COL 1 antibody ((Rabbit polyclonal anti-type I collagen, 600-401-103-0.1 (Rockland) and secondary antibody (goat anti-Rabbit IgG (H+L) Highly Cross-Adsorbed Secondary Antibody, Alexa Fluor 647, A-21245 (dilution 1:200) to visualize H_47_-bound to collagen.

For immunostaining, the cultures were decellularized using a previously established protocol [75], fixed using 4% PFA and stained with Collagen-specific antibodies. Cultures were treated with 0.5% Triton X-100 and 20mM NH_4_OH for 5 min at 37 °C for decellularization. The deposited matrix on the culture substrate was blocked with 5% Goat serum in PBS and stained for COL I, III, IV, V and XII with primary antibodies as recommended by supplier (Rabbit polyclonal anti-type I collagen, 600-401-103-0.1 (Rockland), Collagen III Polyclonal Antibody (Thermo fisher, PA5-34787), Anti-COL4A3 antibody (Sigma, HPA042064-100UL), Anti-Collagen V antibody (ab7046), Anti-COL12A1 antibody (Sigma, HPA070695) (dilution 1:200 for all the antibodies)). Samples were washed 3 times with PBS and stained with secondary antibody (goat anti-Rabbit IgG (H+L) Highly Cross-Adsorbed Secondary Antibody, Alexa Fluor 647, A-21245 (dilution 1:200)). For analysis, the mean gray value of fluorescence intensity was measured for each collagen subtypes and subtracted from background mean gray value of fluorescence intensity by imaging stained collagen with only secondary antibody. Results were normalized by taking untreated MEF cells as 1. In MEF Hsp47 −/− untreated cells were considered as 1. This analysis was performed in Image J.

### Western Blot to study increase in collagen subtypes on treatment with H_47_ in MEF Hsp47 −/− cells

Deposited collagen from cells cultured in the presence and in the absence of H_47_ was suspended in 300 μL of RIPA Buffer. Before this step, the collagen deposited was decellularized using the above-mentioned protocol. Protease inhibitor and Lameli buffer (4x stock concentration) was added in mixture to avoid protein degradation. The samples were loaded into SDS-PAGE gels. The 12% SDS PAGE gels were transferred using blotting chamber to PVDF membranes. The Blotted PVDF Membranes were blocked with Blocking buffer (0.5% milk powder in PBST (0.1 w/v)) for 20 mins. The excess blocking buffer was washed off three times using PBST. The PVDF membranes were incubated overnight at 4°C with: rabbit polyclonal anti-type I collagen (Rockland, 600-401-103-0.1), Collagen III Polyclonal Antibody (Thermo fisher, PA5-34787), Anti-COL4A3 antibody (Sigma, HPA042064-100UL), anti-Collagen V antibody (ab7046), or anti-COL12A1 antibody (Thermo fisher, HPA070695) at 1:200 dilution. For α-Smooth Muscle (α-SMA) condition cells were not decellularized and antibodies used were Anti-Actin, α-Smooth Muscle - Cy3 (C6198) On the following day the excess was washed off three times using PBST (0.5 w/v) and the sample was stained with secondary antibody for 1 h at room temperature (Goat anti-Rabbit IgG (H+L) Highly Cross-Adsorbed Secondary Antibody, Alexa Fluor 647, A-21245 (1:500 dilution). The PVDF membrane after staining was visualized under Gel Doc. All the experiments were done in triplicate.

### Statistical significance

For Sirus Red assay (n=6) number of experiment were performed and plotted with graphs including whisker plots representing standard deviation for MEF, L929, NHDF, HTCAT and HDMEC. For MEF Hsp47 −/− three independent experiments were performed. For immunostaining quantification IMAGE J was used to quantify mean intensity profile of stained matrices in all cell types treated with and without H_47_ with bar plots including error bars indicating standard deviation. In both assays the data were normalized with respect to collagen deposited in MEF untreated as 1. In case of MEF Hsp47 −/− untreated cells were considered as 1. Statistical significance was analyzed by Tukey test, which shows significant differences between conditions. Significance was calculated by comparing non treated vs treated cells (mean±SD, ANOVA, *** p<0.001).

## Additional files

### Additional file 1

Figure S1 shows supplementary information on delivery of H_47_ to ER via KDEL receptor-mediated endocytosis to different cell types.

### Additional file 2

Figure S2 shows Z-stack orthogonal projection images of NHDF after incubation with EGFP for 3 h.

### Additional file 3

Figure S3 shows Immunostaining of COL I, III, IV, V and XII deposited in MEF, L929, HaCaT and HDMEC cultures 24 h with and without treatment of H_47_.

### Additional file 4

Figure S4 shows Stimulated deposition of COL I, III and V in MEF Hsp47 −/− cells after H_47_ uptake.

### Additional file 5

Figure S5 shows H_47_ binds to collagen on the matrix on L929 cells reaching confluency.

## Acknowledgements

The authors thank Prof. Kazuhiro Nagata, Kyoto University (JP) for sharing MEF Hsp47 (+/+) and (−/−) cell lines and Prof. Mathias Laschke, Institut für Klinisch-Experimentelle Chirurgie, UKS, Homburg, (DE) for sharing HDMEC cell line. SK and AdC acknowledge financial support from the Deutsche Forschung Gemeinschaft (SFB 1027). AdC acknowledges funding from the European Union’s Horizon 2020 research and innovation Program No. 731957 (FET Mechanocontrol).

## Author’s contributions

ESK, SS and AdC wrote the manuscript. AdC and SS supervised the study. Experiments were designed by ESK and AdC. Experiments were performed by ESK with assistance from LL. Data quantification and analysis was performed by ESK and interpretation was performed by ESK, SS and AdC.

## Availability of data and materials

All data generated or analyzed during this study are included in this published article and its supplementary information files.

## Ethics approval and consent to participate

Not applicable.

## Consent for publication

Not applicable.

## Additional files

**Figure S1.**
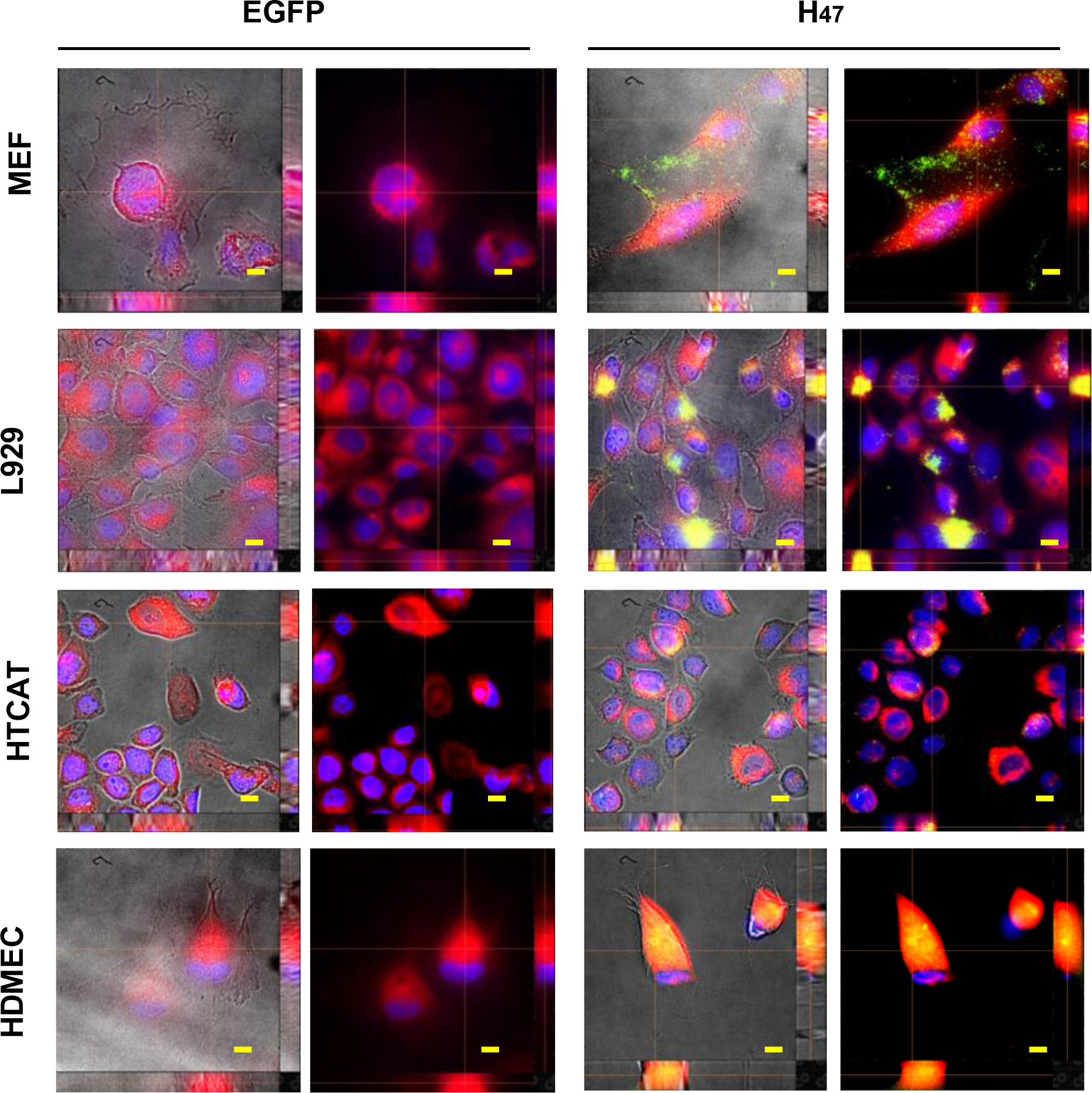
Delivery of H_47_ to ER via KDEL receptor-mediated endocytosis. Z-stack orthogonal projection images of L929, MEF, HaCaT and HDMEC after incubation with H_47_ for 3 h. Images show co localization of H_47_ and ER signals. (Blue: DAPI (Nucleus), Green: (Hsp47), and Red: ER tracker dye). Left side: EGFP treated cell lines. Right side: H_47_ treated cell lines. Scale: 20 μm.

**Figure S2.**
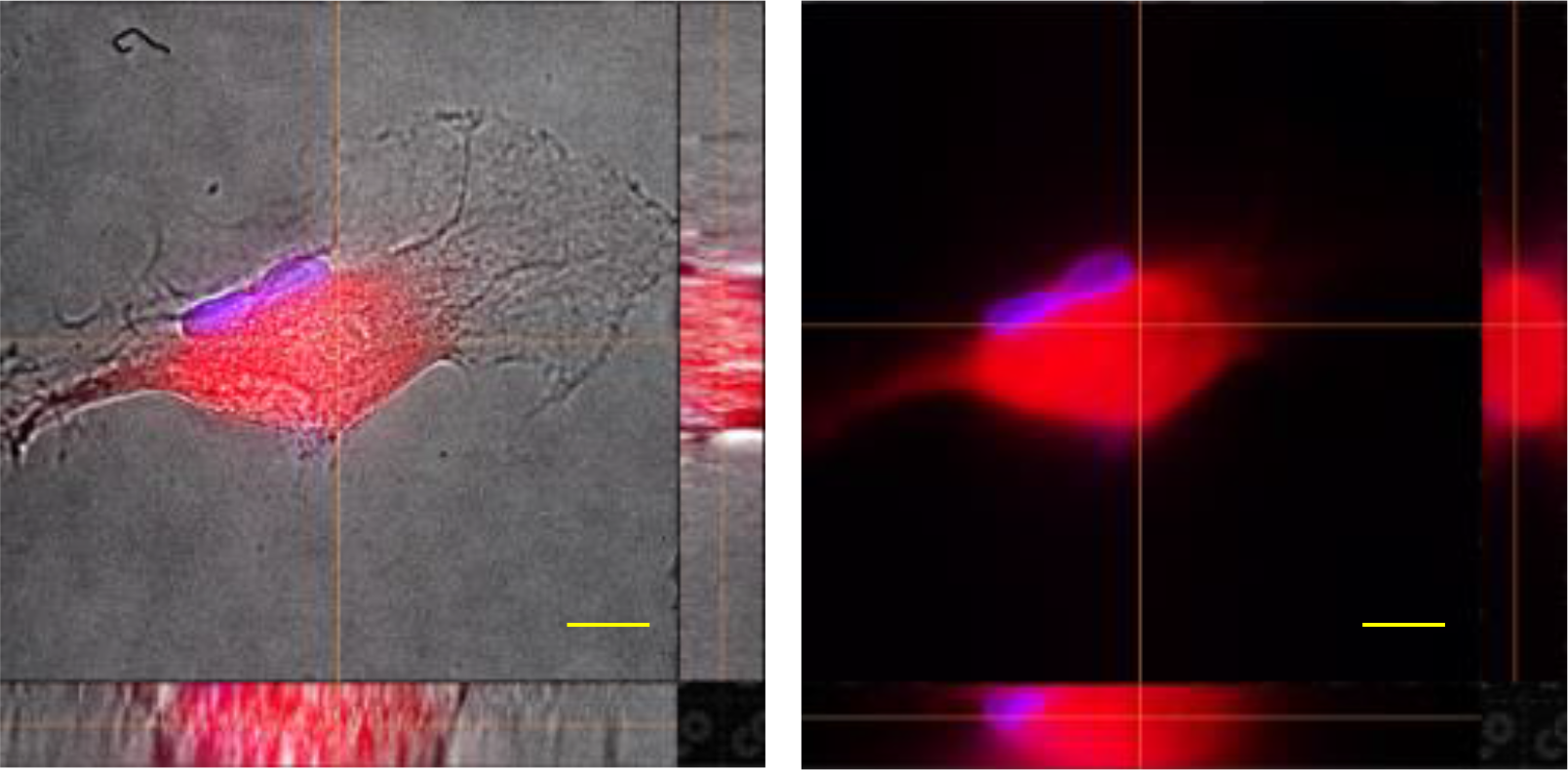
Z-stack orthogonal projection images of NHDF after incubation with EGFP for 3 h. Image show ER signal. (Blue: DAPI (Nucleus), and Red: ER tracker dye). Scale: 20 μm.

**Figure S3.**
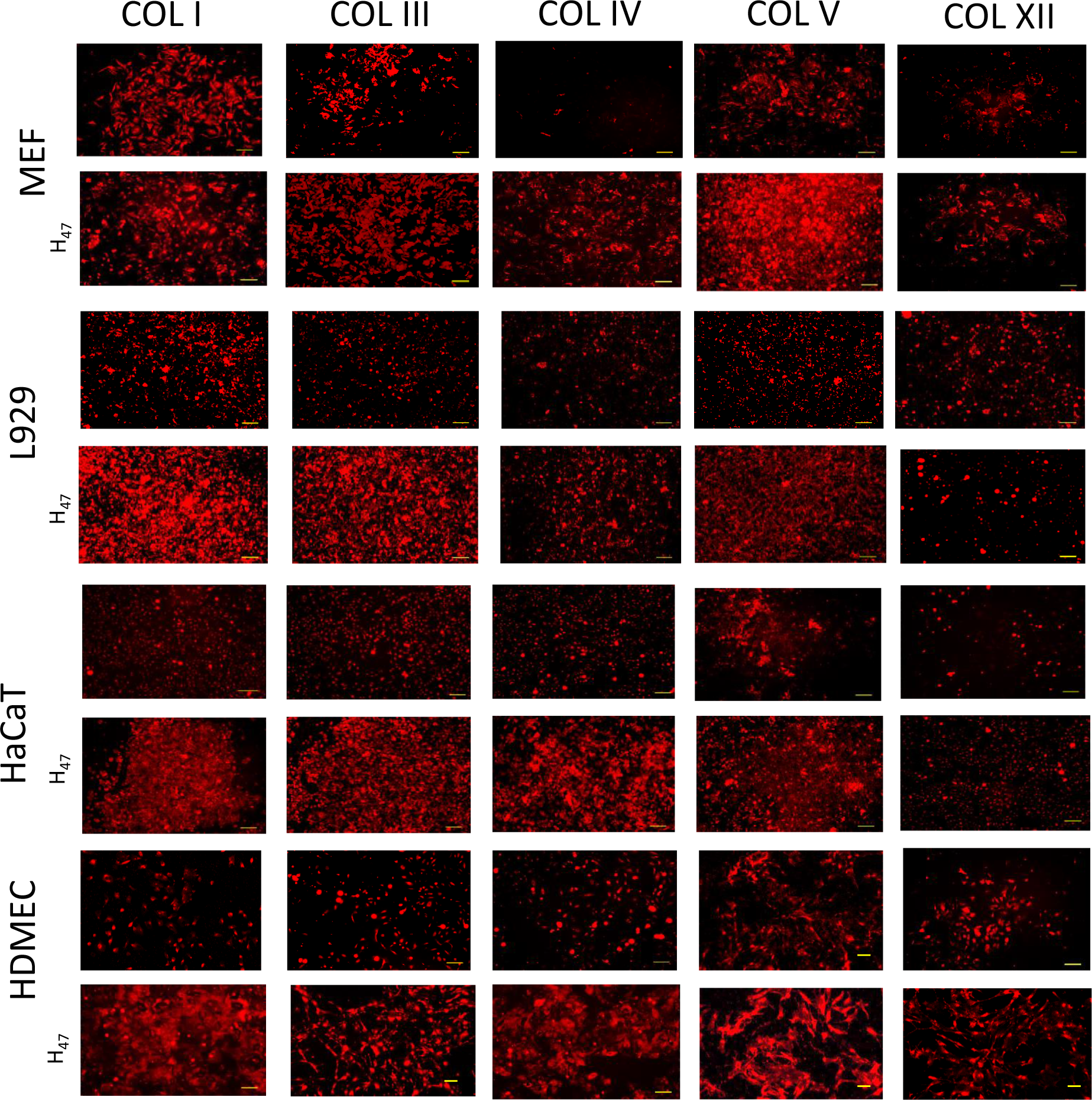
Immunostaining of COL I, III, IV, V and XII. deposited in MEF, L929, HaCaT and HDMEC cultures, 24 h after either no treatment or treatment with H_47_. (Scale corresponds to 250μm).

**Figure S4.**
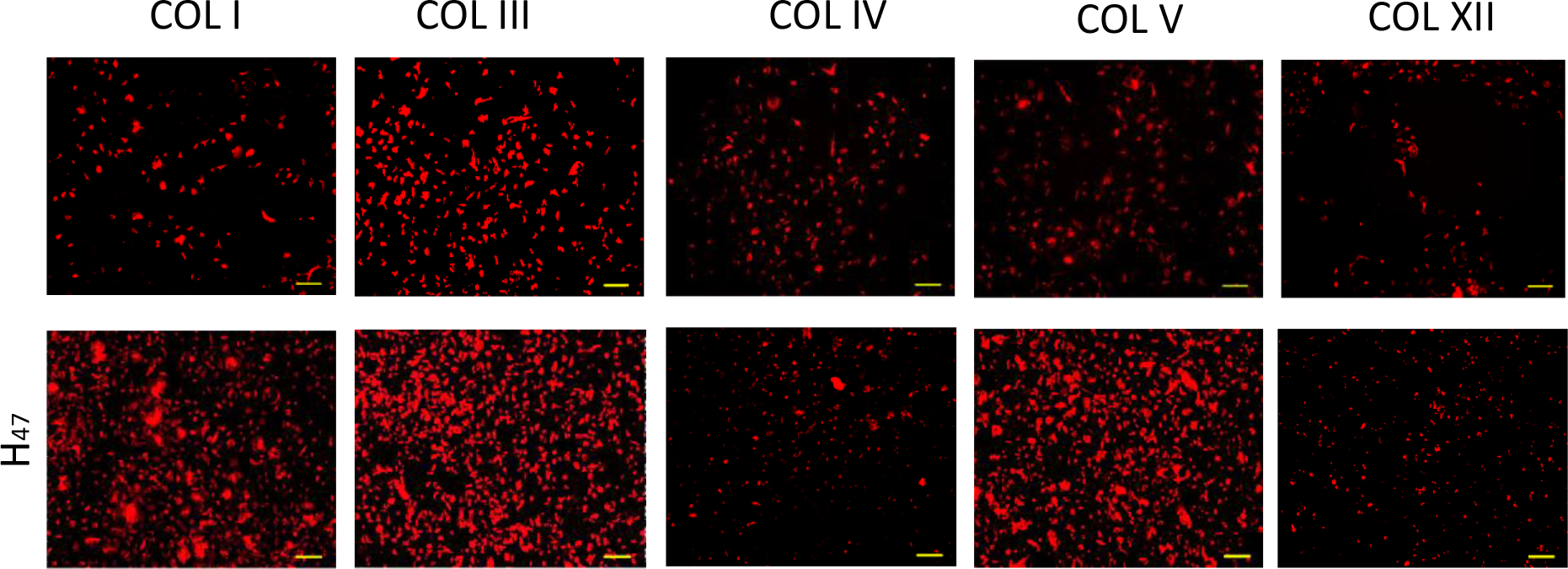
Stimulated deposition of COL I, III and V in MEF Hsp47 −/− cells after H_47_ uptake. **a.**Fluorescence images of immunostained COL I, III, IV, V and XII in MEF Hsp47−/− cells 24 after H_47_ treatment and controls (Scale – 250μm).

**Figure S5.**
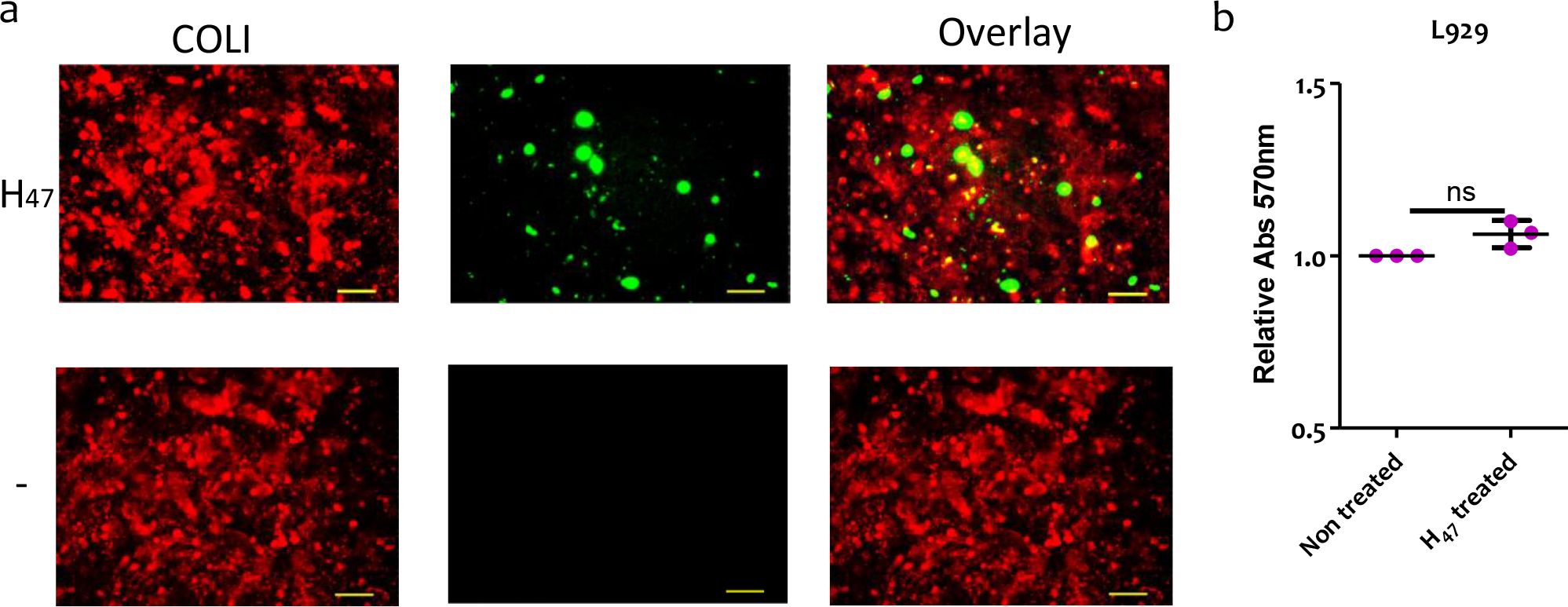
H_47_ binds to collagen on the matrix on L929 cells reaching confluency. a. Fluorescence images of immunostained COL I (Red signal) of confluent L929 cells treated with and without H_47_ (Green signal) (Scale: 250μm). b. Quantification of collagen deposition using Sirius Red assay with error bars representing standard deviation of 3 independent experiments after 24h of H_47_ treatment of confluent L929 cells. The plots in assays were normalized with untreated L929 condition as 1. Statistical significance for was analyzed by t-test test comparing untreated against H_47_ treated conditions (mean±SD, *** p value = 0.0558) (ns=no significance).

## References

1. A.Holzapfel G: Biomechanics of Soft Tissue. In. Edited by Holzapfel GA: BIOMECH PREPRINT SERIES; 2000.

2. Lynne T. Smith KAH, Joseph A. Madri: Collagen Types I, III, and V in Human Embryonic and Fetal Skin.

3. Boerboom RA, Krahn KN, Megens RTA, van Zandvoort, J. Mam, Merkx M, Bouten CVC: High resolution imaging of collagen organisation and synthesis using a versatile collagen specific probe. J Struct Biol 2007, 159(3):392–399.

4. Koppenol DC, Vermolen FJ, Niessen FB, van Zuijlen PPM, Vuik K: A biomechanical mathematical model for the collagen bundle distribution-dependent contraction and subsequent retraction of healing dermal wounds. Biomech Model Mechanobiol 2017, 16(1):345–361.

5. Uhlen M, Fagerberg L, Hallstrom BM, Lindskog C, Oksvold P, Mardinoglu A, Sivertsson A, Kampf C, Sjostedt E, Asplund A et al: Proteomics. Tissue-based map of the human proteome. Science 2015, 347(6220):1260419.

6. Edqvist PH, Fagerberg L, Hallstrom BM, Danielsson A, Edlund K, Uhlen M, Ponten F: Expression of human skin-specific genes defined by transcriptomics and antibody-based profiling. J Histochem Cytochem 2015, 63(2):129–141.

7. Ricard-Blum S: The collagen family. Cold Spring Harb Perspect Biol 2011, 3(1):a004978.

8. Kajbafzadeh AM, Elmi A, Mazaheri P, Talab SS, Jan D: Genitourinary involvement in epidermolysis bullosa: clinical presentations and therapeutic challenges. BJU Int 2010, 106(11):1763–1766.

9. Kuivaniemi H TG, Prockop DJ: Mutations in fibrillar collagens (types I, 11, 111, and XI), fibril-associated collagen (type IX), and network-forming collagen (type X) cause a spectrum of diseases of bone, cartilage, and blood vessels. Hum Mutat 1997, 9:300–315.

10. Myllyharju J, Kivirikko KI: Collagens and collagen-related diseases. Annals of Medicine 2009, 33(1):7–21.

11. Coulombe PA, Kerns ML, Fuchs E: Epidermolysis bullosa simplex: a paradigm for disorders of tissue fragility. J Clin Invest 2009, 119(7):1784–1793.

12. Panwar P, Butler GS, Jamroz A, Azizi P, Overall CM, Brömme D: Aging-associated modifications of collagen affect its degradation by matrix metalloproteinases. Matrix Biology 2018, 65:30–44.

13. Fligiel SEG, Varani J, Datta SC, Kang S, Fisher GJ, Voorhees JJ: Collagen Degradation in Aged/Photodamaged Skin In Vivo and After Exposure to Matrix Metalloproteinase-1 In Vitro. Journal of Investigative Dermatology 2003, 120(5):842–848.

14. Robertson WV, B.: The effect of ascorbic acid deficiency on the collagen concentration of newly induced fibrous tissue. J Biol Chem 1952, 198:403–408.

15. Phillip JM, Aifuwa I, Walston J, Wirtz D: The Mechanobiology of Aging. Annu Rev Biomed Eng 2015, 17:113–141.

16. Pan X, Chen Z, Huang R, Yao Y, Ma G: Transforming growth factor beta1 induces the expression of collagen type I by DNA methylation in cardiac fibroblasts. PLoS One 2013, 8(4):e60335.

17. JC Gessin LB, JS Gordon JS, R. G. Berg: Regulation of collagen synthesis in human dermal fibroblasts in contracted collagen gels by ascorbic acid, growth factors, and inhibitors of lipid peroxidation. Exp Cell Res 1993, 206:283–290.

18. Pullar JM, Carr AC, Vissers MCM: The Roles of Vitamin C in Skin Health. Nutrients 2017, 9(8).

19. S RS, R P, P S, J T, rews: Effect of vitamin C on collagen biosynthesis and degree of birefringence in polarization sensitive optical coherence tomography (PS-OCT). African Journal of Biotechnology 2008, 7(12):2049–2054.

20. S.J Kim JHP, D.H Kim,Y. Ho,H.I. Maibach: Increased In Vivo Collagen Synthesis and In Vitro Cell Proliferative Effect of Glycolic Acid. Dermatologic Surg 1998, 24(10):1054–1058.

21. Besio R, Iula G, Garibaldi N, Cipolla L, Sabbioneda S, Biggiogera M, Marini JC, Rossi A, Forlino A: 4-PBA ameliorates cellular homeostasis in fibroblasts from osteogenesis imperfecta patients by enhancing autophagy and stimulating protein secretion. Biochim Biophys Acta Mol Basis Dis 2018, 1864(5 Pt A):1642–1652.

22. Ganceviciene R, Liakou AI, Theodoridis A, Makrantonaki E, Zouboulis CC: Skin anti-aging strategies. Dermatoendocrinol 2012, 4(3):308–319.

23. Khan ES, Sankaran S, Paez JI, Muth C, Han MKL, Del Campo A: Photoactivatable Hsp47: A Tool to Regulate Collagen Secretion and Assembly. Adv Sci (Weinh) 2019, 6(9):1801982.

24. Byers PH, Pyott SM: Recessively inherited forms of osteogenesis imperfecta. In: Annual Review of Genetics. vol. 46; 2012: 475–497.

25. Dewavrin JY, Abdurrahiem M, Blocki A, Musib M, Piazza F, Raghunath M: Synergistic rate boosting of collagen fibrillogenesis in heterogeneous mixtures of crowding agents. J Phys Chem B 2015, 119(12):4350–4358.

26. Greenspan DS: Biosynthetic Processing of Collagen Molecules. In: Collagen Topics in Current Chemistry. vol. 247; 2005: 149–183.

27. He W, Dai C: Key Fibrogenic Signaling. Current Pathobiology Reports 2015, 3(2):183–192.

28. Makareeva E, Leikin S: Procollagen triple helix assembly: an unconventional chaperone-assisted folding paradigm. PLoS One 2007, 2(10):e1029.

29. Widmer C, Gebauer JM, Brunstein E, Rosenbaum S, Zaucke F, Drogemuller C, Leeb T, Baumann U: Molecular basis for the action of the collagen-specific chaperone Hsp47/SERPINH1 and its structure-specific client recognition. Proc Natl Acad Sci U S A 2012, 109(33):13243–13247.

30. Mamoru Satoh KH, Shin-ichi Yokota, Nobuko Hosokawa, and Kazuhiro Nagata: Intracellular Interaction of Collagen-specific Stress Protein HSP47with Newly Synthesized Procollagen The Journal of Cell Biology 1996, 133:469–483.

31. Ito S, Nagata K: Roles of the endoplasmic reticulum-resident, collagen-specific molecular chaperone Hsp47 in vertebrate cells and human disease. J Biol Chem 2019, 294(6):2133–2141.

32. Koide T, Asada S, Takahara Y, Nishikawa Y, Nagata K, Kitagawa K: Specific recognition of the collagen triple helix by chaperone HSP47: Minimal structural requirement and spatial molecular orientation. Journal of Biological Chemistry 2006, 281(6):3432–3438.

33. Thomson CA, Ananthanarayanan VS: Structure-function studies on hsp47: pH-dependent inhibition of collagen fibril formation in vitro. J biol Chem.

34. Thomson CA, Tenni R, Ananthanarayanan VS: Mapping Hsp47 binding site(s) using CNBr peptides derived from type I and type II collagen. Protein Science 2003, 12(8):1792–1800.

35. Christy A. Thomson VSA: Structure–function studies on Hsp47: pH-dependent inhibition of collagen fibril formation in vitro. Biochem J 2000, 349(877–883).

36. Thomson CA, Tenni R, Ananthanarayanan VS: Mapping Hsp47 binding site(s) using CNBr peptides derived from type I and type II collagen. Protein Sci 2003, 12(8):1792–1800.

37. Razzaque M: Expression profiles of collagens, HSP47, TGF-β1, MMPs and TIMPs in epidermolysis bullosa acquisita. Cytokine 2003, 21(5):207–213.

38. Zhu J, Xiong G, Fu H, Evers BM, Zhou BP, Xu R: Chaperone Hsp47 Drives Malignant Growth and Invasion by Modulating an ECM Gene Network. Cancer Research 2015, 75(8):1580–1591.

39. Yamamoto N, Kinoshita T, Nohata N, Yoshino H, Itesako T, Fujimura L, Mitsuhashi A, Usui H, Enokida H, Nakagawa M et al: Tumor-suppressive microRNA-29a inhibits cancer cell migration and invasion via targeting HSP47 in cervical squamous cell carcinoma. International Journal of Oncology 2013, 43(6):1855–1863.

40. Duarte BDP, Bonatto D: The heat shock protein 47 as a potential biomarker and a therapeutic agent in cancer research. Journal of Cancer Research and Clinical Oncology 2018, 144(12):2319–2328.

41. Canty EG, Kadler KE: Procollagen trafficking, processing and fibrillogenesis. J Cell Sci 2005, 118(Pt 7):1341–1353.

42. Ishikawa Y, Bachinger HP: A molecular ensemble in the rER for procollagen maturation. Biochim Biophys Acta 2013, 1833(11):2479–2491.

43. Masago Y, Hosoya A, Kawasaki K, Kawano S, Nasu A, Toguchida J, Fujita K, Nakamura H, Kondoh G, Nagata K: The molecular chaperone Hsp47 is essential for cartilage and endochondral bone formation. J Cell Sci 2012, 125(Pt 5):1118–1128.

44. Nakai A: Involvement of the stress protein HSP47 in procollagen processing in the endoplasmic reticulum. The Journal of Cell Biology 1992, 117(4):903–914.

45. Mirco Capitani, Sallese M: The KDEL receptor: New functions for an old protein. FEBS Letters 2009:3863–3871.

46. Becker B, Shaebani MR, Rammo D, Bubel T, Santen L, Schmitt MJ: Cargo binding promotes KDEL receptor clustering at the mammalian cell surface. Sci Rep 2016, 6:28940.

47. Cabrera M, Muniz M, Hidalgo J, Vega L, Martin ME, Velasco A: The retrieval function of the KDEL receptor requires PKA phosphorylation of its C-terminus. Mol Biol Cell 2003, 14(10):4114–4125.

48. Tomohiko Aoe AJL, Elly van Donselaar, Peter J. Peters, and Victor W. Hsu: Modulation of intracellular transport by transported proteins: Insight from regulation of COPI-mediated transport. Proc Natl Acad Sci 1998, 95:1624–1629.

49. Ito S, Ogawa K, Takeuchi K, Takagi M, Yoshida M, Hirokawa T, Hirayama S, Shin-Ya K, Shimada I, Doi T et al: A small-molecule compound inhibits a collagen-specific molecular chaperone and could represent a potential remedy for fibrosis. J Biol Chem 2017, 292(49):20076–20085.

50. Corrine R. Kliment JME, Lauren P. Crum, Tim D. Oury: A novel method for accurate collagen and biochemical assessment of pulmonary tissue utilizing one animal. Int J Clin Exp Pathol 2011, 4(4):349–355.

51. Cole MA, Quan T, Voorhees JJ, Fisher GJ: Extracellular matrix regulation of fibroblast function: redefining our perspective on skin aging. J Cell Commun Signal 2018, 12(1):35–43.

52. Fisher GJ, Varani J, Voorhees JJ: Looking older: fibroblast collapse and therapeutic implications. Arch Dermatol 2008, 144(5):666–672.

53. Mitra M, Ho LD, Coller HA: An In Vitro Model of Cellular Quiescence in Primary Human Dermal Fibroblasts. In: Cellular Quiescence. 2018: 27–47.

54. Mamoru Satoh KH, Shin-ichi Yokota, Nobuko Hosokawa, and Kazuhiro Nagata: Intracellular Interaction of Collagen-specific Stress Protein HSP47 with Newly Synthesized Procollagen. The Journal of Cell Biology 1996, 133:469–483.

55. Elena Dellambra GPD: Cellular Senescence and Skin Aging. In: Skin Aging Handbook: An Integrated Approach to Biochemistry and Product Development. Edited by Dayan N: William Andrew Inc.; 2009: 129–148.

56. Maresa Wick CB, Sabine Briisselbach, Frances C. Lucibello, and Rolf Muller: A Novel Member of Human Tissue Inhibitor of Metalloproteinases (TIMP) Gene Family Is Regulated during G, Progression, Mitogenic Stimulation, Differentiation, and Senescenc. JBiolChem 1994, 269:18953–18960.

57. Dawne N. Shelton EC, Peter S. Whittier, Donghee Choi and Walter D. Funk: Microarray analysis of replicative senescence. Current Biology 1999, 9.

58. Aumailley M KT, Razaka G, Müller PK, Bricaud H.: Influence of cell density on collagen biosynthesis in fibroblast cultures. Biochem J 1982, 206(3):505–510.

59. Ishida Y, Kubota H, Yamamoto A, Kitamura A, Bachinger HP, Nagata K: Type I collagen in Hsp47-null cells is aggregated in endoplasmic reticulum and deficient in N-propeptide processing and fibrillogenesis. Mol Biol Cell 2006, 17(5):2346–2355.

60. Ishida Y, Nagata K: Hsp47 as a collagen-specific molecular chaperone. In: Methods in Enzymology. vol. 499; 2011: 167–182.

61. Nagai N, Hosokawa M, Itohara S, Adachi E, Matsushita T, Hosokawa N, Nagata K: Embryonic Lethality of Molecular Chaperone Hsp47 Knockout Mice Is Associated with Defects in Collagen Biosynthesis. The Journal of Cell Biology 2000, 150(6):1499–1506.

62. Kuroda K, Tsukifuji R, Shinkai H: Increased expression of heat-shock protein 47 is associated with overproduction of type I procollagen in systemic sclerosis skin fibroblasts. J Invest Dermatol 1998, 111(6):1023–1028.

63. Wang ZL, Inokuchi T, Ikeda H, Baba TT, Uehara M, Kamasaki N, Sano K, Nemoto TK, Taguchi T: Collagen-binding heat shock protein HSP47 expression during healing of fetal skin wounds. Int J Oral Maxillofac Surg 2002, 31(2):179–184.

64. Johnston EF, Gillis TE: Transforming growth factor beta-1 (TGF-beta1) stimulates collagen synthesis in cultured rainbow trout cardiac fibroblasts. J Exp Biol 2017, 220(Pt 14):2645–2653.

65. Lu Y, Azad N, Wang L, Iyer AK, Castranova V, Jiang BH, Rojanasakul Y: Phosphatidylinositol-3-kinase/akt regulates bleomycin-induced fibroblast proliferation and collagen production. Am J Respir Cell Mol Biol 2010, 42(4):432—441.

66. Liu Y, Li Y, Li N, Teng W, Wang M, Zhang Y, Xiao Z: TGF-beta1 promotes scar fibroblasts proliferation and transdifferentiation via up-regulating MicroRNA-21. Sci Rep 2016, 6:32231.

67. Moses DHL: TGF-P Regulation of Epithelial Cell Proliferation. Mol Reprod Dev 1992, 32:179–184.

68. Trompezinski S, Berthier‐Vergnes, O., Denis, A., Schmitt, D. and Viac, J: Comparative expression of vascular endothelial growth factor family members, VEGF‐B, ‐C and ‐D, by normal human keratinocytes and fibroblasts. Experimental Dermatology 2004, 13:98–105.

69. Jeffrey M. Davidson PAL, Ornella Zoia‡, Daniela Quaglino, Jr.,MariaGabriella Giro: Ascorbate Differentially Regulates Elastin and Collagen Biosynthesis in Vascular Smooth Muscle Cells and Skin Fibroblasts by Pretranslational Mechanisms. JBiolChem 1997, 272:345–352.

70. Murad S, Tajima S, Johnson GR, Sivarajah A, Pinnell SR: Collagen Synthesis in Cultured Human Skin Fibroblasts: Effect of Ascorbic Acid and Its Analogs. Journal of Investigative Dermatology 1983, 81(2):158–162.

71. Seong-Jin Kim. Jong-Jin Kim J-HP, Do-Heong-Jin Kim,Jong-Hyuk Park, Do-Hyun Kim, Young-Ho Won, Howard I. Maibach Increased In Vivo Collagen Synthesis and In Vitro Cell Proliferative Effect of Glycolic Acid. Dermatol Surg 1998, 24:1054–1051 1058.

72. Varani; Gary J. Fisher; John J. Voorhees; Sewon Kang RKHSRKWESSCVNHTAHALKJDNJ: Improvement of Naturally Aged Skin With Vitamin A (Retinol). ARCH DERMATOL 2007, 143:606–612.

73. Sevilla CA, Dalecki D, Hocking DC: Regional fibronectin and collagen fibril co-assembly directs cell proliferation and microtissue morphology. PLoS One 2013, 8(10):e77316.

74. Duval K, Grover H, Han LH, Mou Y, Pegoraro AF, Fredberg J, Chen Z: Modeling Physiological Events in 2D vs. 3D Cell Culture. Physiology (Bethesda) 2017, 32(4):266–277.

75. Pinto ML, Rios E, Silva AC, Neves SC, Caires HR, Pinto AT, Duraes C, Carvalho FA, Cardoso AP, Santos NC et al: Decellularized human colorectal cancer matrices polarize macrophages towards an anti-inflammatory phenotype promoting cancer cell invasion via CCL18. Biomaterials 2017, 124:211–224.

